# rAmpSeq: Using repetitive sequences for robust genotyping

**DOI:** 10.1101/096628

**Authors:** Edward S. Buckler, Daniel C. Ilut, Xiaoyun Wang, Tobias Kretzschmar, Michael Gore, Sharon E. Mitchell

## Abstract

Repetitive sequences have been used for DNA fingerprinting and genotyping for more than a quarter century. Now, with our knowledge of whole genome sequences, repetitive sequences can be used to identify polymorphisms that can be mapped and scored in a systematic manner. We have developed a simple, robust platform for designing primers, PCR amplification, and high throughput cloning that allows hundreds to thousands of markers to be scored for less than $5 per sample. Conserved regions were used to design PCR primers for amplifying thousands of middle repetitive regions of the maize (*Zea mays* ssp. *mays*) genome. Bioinformatic scans were then used to identify DNA sequence polymorphisms in the low copy intervening sequences. When used in conjunction with simple DNA preps, optimized PCR conditions, high multiplex Illumina indexing and a bioinformatic marker calling platform tailored for repetitive sequences, this methodology provides a cost effective genotyping strategy for large-scale genomic selection projects. We show detailed results from four maize primer sets that produced between 1,335-3,225 good coverage loci with 1056 that segregated appropriately in a bi-parental family. This approach could have wide applicability to breeding and conservation biology, where hundreds of thousands of samples need to be genotyped for very minimal cost.

## Introduction

Genomics has revolutionized our understanding of the genetic architecture of organisms, evolution, and molecular mechanisms. On the applied side, the simple use of genomic profiling of natural variation has revolutionized animal and plant breeding in the last decade [1,2]. Genomic selection, a type of marker-assisted breeding where all quantitative trait loci (QTL) are in linkage disequilibrium with genome-wide markers, has been most successfully applied to organisms that are expensive to phenotype (e.g. cattle) or where custom genotyping platforms could drive accelerated breeding cycles (e.g. maize). Despite this progress, genomic selection’s greatest gains are likely still to come. There are tremendous opportunities to apply genomic selection to thousands of animal and crop efforts, forests trees, and aquaculture systems, where relatively little selection and modern breeding techniques have not been applied. Similar opportunities are available to conservation biology, where thousands of organisms need to be genotyped, but the resources are rarely available for a focused effort in any one species.

The goal of this research was to develop a genotyping assay that could be applied to most species, use low quality DNA, generate several hundred to a few thousand markers that would facilitate whole genome imputation, and eventually cost less than $2 per sample. Most of the current technologies do not provide this level of performance and adaptability to diverse molecular lab facilities, where the next generation of species are being studied. To create a robust, simple protocol, we combine the 30 year-old concepts of PCR-amplifying and genotyping repetitive sequences with next-generation sequencing to make a simple assay.

Restriction enzyme-based genotyping-by-sequencing (GBS) has had tremendous impact over the last 16 years [3–6]. The- key impact was providing a reduced representation library that focuses sequencing on a modest portion of the genome, and these approaches could be applied to any species without any prior knowledge of the genome. However, there are two main limitations that have been observed with restriction enzyme-mediated GBS:

1. High quality DNA is generally needed to efficiently use restriction enzymes, which adds substantially to the cost, time, and effort necessary for the entire genotyping process. This has been a serious obstacle in plant breeding, where 100,000s of samples need to be processed rapidly. To address the DNA quality issue, we have shifted the first step of the genotyping assay to standard, exponential PCR. The amplification conditions and polymerases used in PCR have become quite robust to DNA contaminants and have always been very sensitive.
2. Whenever PCR encounters thousands of heterogenous amplicons, there is PCR competition between the various amplicons. Amplicon lengths, GC contents, and hairpin structures all combine to determine the efficiency of amplification. In the case of restriction GBS, the PCR competition had the positive benefit of naturally providing size selection. However, competition also produces unequal coverage between various amplicons, which results in a large proportion of scoring dropouts. These dropouts can make scoring heterozygous loci more difficult. Linear amplification provides a mechanism to reduce this variability [7], however, another option is to focus on the naturally repetitive sequences in a genome. By targeting the repetitive fraction of the genome, the sequences are nearly identical in length and composition, which reduces competition issues.

Amplicon sequencing has become an alternative that is certainly very capable of simultaneously genotyping 10s to 100s of markers. While amplicon sequencing approaches are nearly as old as PCR, the throughput of next-generation sequencers is allowing these methods to become quite cost effective, e.g. AmpSeq [8]. The limitation of this approach is generally the prior knowledge of numerous SNPs, essentially a HapMap, and there can be amplicon bias and primer competition that prevents a few thousand markers from being multiplexed without extensive optimization. In this study, we focus amplicon sequencing at the repetitive fraction of the genome - essentially AmpSeq with one or few primer pairs.

Genotyping the repetitive fraction of a genome is actually one of the oldest approaches to genotyping, as initially these regions were well characterized and were easy to clone. Ribosomal sequences have been a target for studying inter- and intra-species phylogenetics for three decades [9]. The key design for genotyping-by-sequencing with ribosomal sequences was designing primers on the conserved regions of the ribosomal sequences (26S, 5.8S, and 18S) that span the less conserved internal transcribed spacers (ITS). Intra-individual and intra-populational variation among thousands of ribosomal repeats was scored using Sanger sequencing of these regions [10–12], and used for evolutionary and population genetics. Similarly, in human genetics the repetitive MHC loci have have been genotyped through PCR amplification, cloning, and sequencing [13]. The sequencing of repetitive sequences has also given rise to the entire field of metagenomics, where ribosomal repeat sequencing is used to identify the trillions of organisms present everywhere.

For the last three decades, it has been clear that transposons dominate the genomes of most higher eukaryotes. Once this was realized, transposable elements became targets for genotyping [14]. Transposon display moved this type of genotyping to repetitive elements that were evenly spread across the genome [15]. Genotyping by scoring random regions between the conserved elements of transposons was also used [16].

The main limitation of using repetitive sequences for intraspecific genotyping has been that the length and quality of DNA sequencing needs to be sufficient to differentiate closely homologous sequences. This was possible with Sanger sequencing by 1990, and then long read pyrosequencing, but until recently sequencing by synthesis (SBS) approaches did not provide the length and quality necessary to differentiate closely repetitive sequences.

Modern SBS sequencers have essentially solved this problem, where high quality 150bp sequences, with or without pairedend reads, can clearly differentiate among close homologs. While repetitive sequences are highly homologous, it is common to find families that are only 80-90% identical for portions of the sequences. With 150 bp reads, 15 to 30 single nucleotide changes that differentiate a pair of paralogs may be obtained.

Here, we combine novel bioinformatics to identify genotyping targets and marker scoring with standard amplification and Illumina-based sequencing to generate a robust genotyping platform that we call rAmpSeq for <u>r</u>epeat <u>Amp</u>lification <u>Seq</u>uencing.

We have designed the assay in a way to address the problems with restriction enzyme-based GBS. The design of rAmpSeq does sacrifice three of the strengths of GBS: (1) It will produce fewer markers in general - often on an order of magnitude less. (2) Knowledge of the reference genome sequence is necessary to design a quality assay. (3) GBS markers were frequently focused near genes, while rAmpSeq generally is targeting intergenic regions. We believe these are reasonable sacrifices as long read sequencing technologies have advanced tremendously so that numerous species have quality genomes. Second, low coverage whole genome sequencing is becoming very inexpensive - on the order $50-250 per genome. When combined with imputation, however, only a small proportion of samples need to be sequenced at high coverage.

We evaluate the power of rAmpSeq by genotyping maize, where we demonstrate primer design approaches and considerations, evaluate sensitivity to DNA concentration and quality, library coverage and distribution, and then genotyping with both orthologs and tightly linked paralogs. Finally, we discuss how this methodology can become a highly distributed protocol that can cost $2 per sample and is capable of delivering the genomics revolution to all corners of breeding and conservation biology, including resource and infrastructure-limited institutions in the developing world.

## Methods and Results

### Plant materials and DNA extraction

PCR optimization and preliminary sequencing were performed using maize inbred lines B73, CML247, W22 and Mo17. Further genotyping was then performed on the parents and 48 progeny from a recombinant inbred population, (B73 X CML 247)F_6_ [17] and 48 diverse maize inbreds from the USDA-ARS National Plant Germplasm System (Table 1). DNAs from lyophilized leaf tissues were extracted, by using either a column-based (DNeasy, QIAGEN, Inc; Valencia, CA) or a standard CTAB protocol [18].

**Table 1.** Diverse maize inbred lines.

| Accession Name | GRIN* Identifier | Accession Name | GRIN Identifier | Accession Name | GRIN Identifier |
| --- | --- | --- | --- | --- | --- |
| A188 | Ames 22443 | DE2 | PI 606330 | NC290A | Ames 27141 |
| A239 | Ames 23405 | EP1 | Ames 27111 | NC320 | Ames 27156 |
| A441-5 | Ames 27064 | F6 | NSL 22894 | NC356 | Ames 27174 |
| A654 | PI 587141 | I137TN | Ames 27116 | ND246 | PI 550490 |
| B77 | PI 608765 | I29 | Ames 27115 | Oh43E | Ames 27183 |
| CH701-30 | Ames 27069 | Ky21 | Ames 27130 | Oh603 | PI 573098 |
| CI 7 | Ames 28367 | L317 | NSL 65873 | P39 | Ames 28186 |
| CI 3A | Ames 26116 | M14 | NSL 30867 | R109B | NSL 242484 |
| CI 64 | Ames 19314 | Mo17 | PI 648430 | R168 | NSL 30891 |
| CM174 | Ames 19316 | Mo18W | PI 550441 | SC 213R | NSL 53083 |
| CML 108 | Ames 27082 | Mo47 | PI 583352 | SG 18 | NSL 75979 |
| CML 220 | Ames 27087 | MS1334 | Ames 24752 | Tzi 25 | PI 506255 |
| CML 228 | Ames 27088 | N6 | Ames 27137 | U267Y | Ames 27191 |
| CML 281 | Ames 27092 | NC230 | Ames 12731 | Va35 | PI 587150 |
| CML 287 | PI 595565 | NC232 | Ames 12732 | W22 | PI 674445 |
| CO106 | Ames 27105 | NC250 | PI 550555 | WD | Ames 27195 |

### Primer design

Using the genome indexing approach integrated in the TASSEL pipeline (version 5; http://www.maizegenetics.net/tassel; [19]), we catalogued all 20 bp kmers with 1000 to 2000 copies and selected kmer pairs with at least 90% similarity in GC content (35%< GC < 65%), 3bp of GC-clamp on the 3′ end, and separation of 125 to 200 bp. The kmers were also selected for relatively even distribution across the genome by testing for presence in 40 genomic bins. The top 5000 of these potential primer pairs were ranked such that standard deviation in amplicon length was minimized and sequence divergence between amplicons was maximized. These 5000 primer pairs were then scored with Primer 3.0 [20], and the top 10 primer pairs coming from four different repeat classes were selected for further evaluation (Table 2).

**Table 2.**
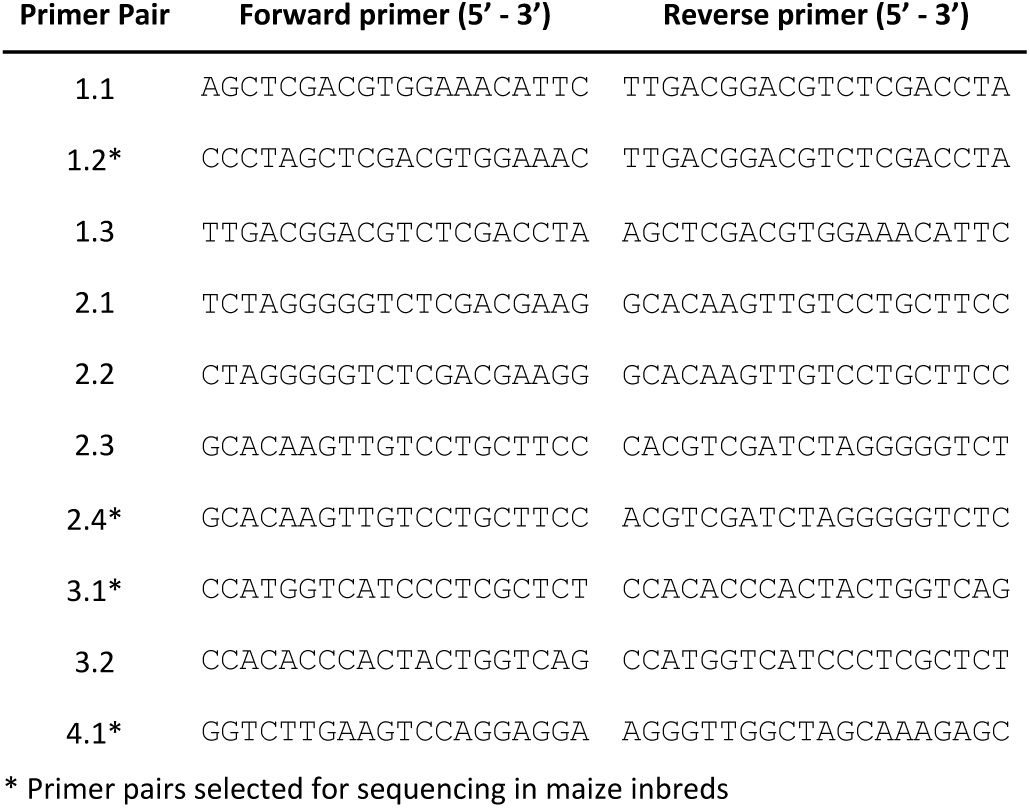
rAmpSeq primer pairs evaluated.

### PCR optimization

Preliminary PCRs were performed in 25μL volumes containing 100 ng genomic DNA (from each of four maize inbred lines), 1X Taq master mix (New England Biolabs, Inc., Ipswich, MA) and 10 pmol of each primer. Standard PCR cycling conditions consisted of 1X 98°C/30 s; 25X (98°C/20 s, 56°C/30 s, 72°C/30 s); followed by a final cycle at 72°C/5 min. A modified cycling protocol designed to reduce competition between PCR artifacts and the desired amplicons, (98°C/30 s; 2X (98°C/20 s, 56°C/30 s, 72°C/30 s), 12X (98°C/20 s, 58°C/30 s, 72°C/30 s); 1X (98°C/20 s, 58°C/2 min with 1°C increase/5 sec, 72°C/30 s), 1 cycle 72°C/5 min) was also evaluated.

To test the limits of the protocol, 15 to 25 cycle standard PCRs were performed with varying amounts of DNA template (five-fold dilutions ranging from 100 ng to 120 picograms of high quality DNA and 100 ng of “poor quality” DNA). Amplification products were purified on QIAquick PCR purification columns according to the manufacturer′s instructions (QIAGEN, Inc., Valencia, CA), eluted in 25 μL volumes and assayed (2μL) on an Experion™ automated electrophoresis station using the Experion 15–1,500 bp DNA Analysis Kit (Bio-Rad Laboratories, Inc., Hercules, CA). Gel images were automatically processed using the included instrument software.

Initial results indicated that the ten rAmpSeq primer pairs amplified robustly, producing measurable products from small amounts (a few hundred picograms) of purified input DNA and degraded, unpurified DNA samples (Figure 1A). All primers produced a band of the predicted size, based on the reference sequence, along with slightly larger (< 50 bp) amplification products. The higher molecular weight fragments, likely the result of competition between correctly primed sites and PCR artifacts, were minimized by reducing the number of PCR cycles, slightly increasing the annealing temperature and using a slow “up ramp” from annealing to the PCR extension temperature in the last PCR cycle (Figure 1B).

**Figure 1.**
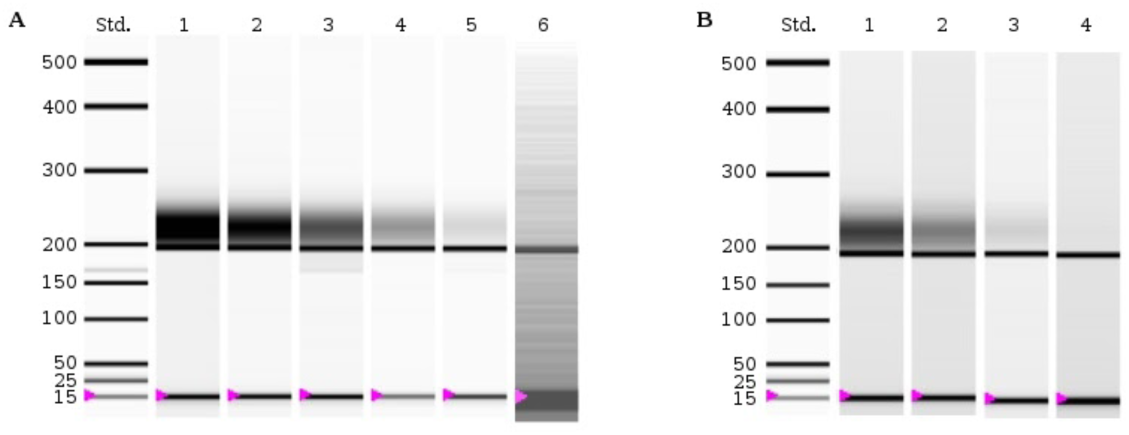
Template titration and PCR effects for rAmpSeq locus 4.1. Template titration experiment (A) and the effect of PCR cycle number and slow annealing on PCR competition (B In A, lanes 1-5 contain amplification products from standard, 20 cycle PCRs using 100 ng, 20 ng, 4 ng, 800 pg and 160 pg DNA from accession B73 as template, respectively. Lane 6 contains amplification products from a degraded, unpurified DNA sample (100 ng) from a standard, 25 cycle PCR. In B, 100 ng of DNA from accession B7 was used as template in all PCRs. Lane 1, standard 20 cycle PCR; Lane 2, 20 cycle PCR with slow primer annealin step; Lane 3, standard 15 cycle PCR; Lane 4, 15 cycle PCR with slow primer annealing step. In all cases, the expected product size is approximately 196 bp. Pink triangles mark the position of the 15 bp size standards, both the ladder and internal standards added to each DNA sample.

### Barcoding and DNA sequencing

Loci 1.2, 2.4, 3.1 and 4.1 (Table 2), were selected for tailed amplicon sequencing in four maize inbred lines (16- plex). After initial results were evaluated, sequencing was repeated in a larger population (n=96) comprising both recombinant inbred lines and a panel of diverse inbreds (384-plex). For compatibility with the Nextera indexing system (Illumina, Inc., San Diego, CA), additional sequences were added to the 5′ ends of the forward (TCGTCGGCAGCGTCAGATGTGTATAAGAGACAG) and reverse (GTCTCGTGGGCTCGGAGATGTGTATAAGAGACAG) primers for each locus. After performing the initial PCRs, as in the modified PCR protocol above, dual indexes were added to the individual amplification products according to the Nextera kit manufacturer’s instructions (Illumina, Inc., San Diego, CA). DNAs (5μL each) from individual reactions were pooled, cleaned using polyethylene glycol-coated magnetic beads (Agilent Technologies, Inc., Santa Clara, CA), quantified and assayed on a BioAnalyzer (Agilent Technologies, Inc., Santa Clara, CA). The four 96-plex libraries were then combined in equal concentrations (2 nM each) and sequences (paired-end, 250 nucleotides) were collected on a MiSeq instrument using version 2 chemistry (RTA version 1.18.54; Illumina, Inc., San Diego, CA). To increase library complexity, phiX DNA (5%) was added to both the 16- and 384-plex libraries prior to sequencing. Sequencing results were converted to FASTQ format, evaluated for overall sequencing quality, and demultiplexed using Illumina bcl2fastq2 software (version 2.17; Illumina, Inc., San Diego, CA).

### Sequence processing

Demultiplexed, barcode-free sequences (reads) were processed using a custom pipeline. The code used in this and all subsequent analysis is available in a public repository (URL:; DOI:). In brief, the raw forward reads for each sample were randomly downsampled to at most 200,000 reads in order to simulate a typical rAmpSeq run, and all sequences that did not pass Illumina quality control were removed. The remaining sequences were trimmed to the first 150 bp, and reverse Illumina Nextera sequences(CTGTCTCTTATACACATCTCCGAGCCCACGAGAC) were removed usingCutadapt(version1.10;https://github.com/marcelm/cutadapt;[21]) with the default 10% maximum mismatch tolerance. Cutadapt was further used to remove the 5′ PCR primer sequence (20% maximum mismatch tolerance, 5′ anchored sequence) and any 3′ sequence that followed the 3′ PCR primer sequence, including the primer sequence itself (no mismatch allowed, unanchored 3′ sequence). Only sequences where the expected 5′ PCR primer sequence was found were retained for further analysis, with both pairs required to pass this filter in the paired end analysis. Finally, low quality bases were removed from the 3′ end using a quality value threshold of 25, and any sequences shorter than 100 bp were discarded.

For each primer pair and sample combination we retained an average of 87% of the reads after filtering, and we were able to identify the expected PCR primer sequence in over 99% of the reads that passed Illumina quality control (Table 3). Detailed statistics of the filtering process are available as supplemental material (Table S2).

**Table 3.**
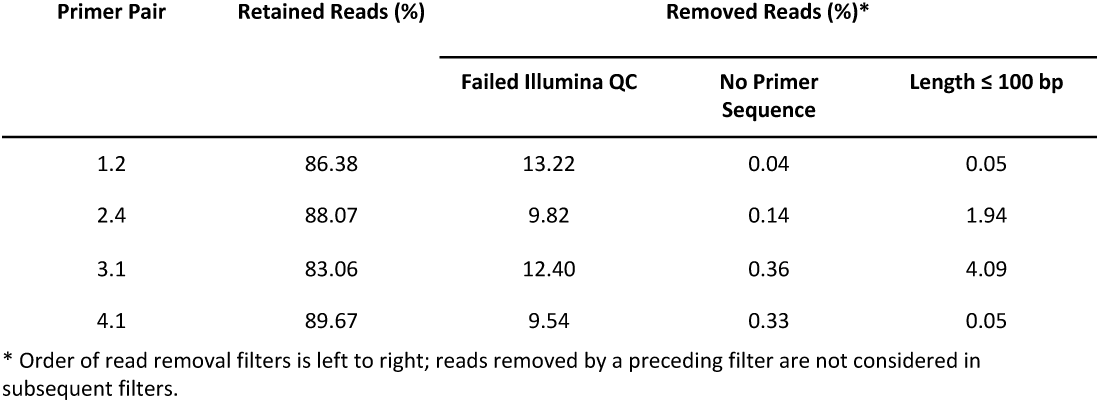
Read filtering results, average values.

| Primer Pair | Retained Reads (%) | Removed Reads (%)* |  |  |
| --- | --- | --- | --- | --- |
|  |  | Failed Illumina QC | No Primer Sequence | Length ≤ 100 bp |
| 1.2 | 86.38 | 13.22 | 0.04 | 0.05 |
| 2.4 | 88.07 | 9.82 | 0.14 | 1.94 |
| 3.1 | 83.06 | 12.40 | 0.36 | 4.09 |
| 4.1 | 89.67 | 9.54 | 0.33 | 0.05 |

\* Order of read removal filters is left to right; reads removed by a preceding filter are not considered in subsequent filters.

### Alignment to the reference genome

Reads from the B73 sample were aligned against the maize reference genome (AGP v3, EnsemblGenomes release 31;[22]) using bwa-mem (version 0.7.12-r1039;https://github.com/lh3/bwa; [23]) with the following parameters: -M -c 100000 -L 5,5 -T 0. The resulting alignment file was further filtered using SAMtools (version 1.3.1; https://github.com/samtools/; [24]), and tags having primary alignments (SAMtools flag -F 0xf0c) with no mismatches on the ten maize chromosomes were retained. They were further filtered to remove those with ambiguous alignments, define as having a secondary alignment with the same alignment score as the primary alignment. Alignment results were processed with BEDTools (version 2.25.0; https://github.com/arq5x/bedtools2; [25]) to merge overlapping alignment locations on the same strand and calculate read depth and average alignment quality for each alignment locus in the genome.

We estimated the number of loci amplified by each sequenced primer pair by aligning the reads from the B73 sample against the maize reference genome, which is constructed from the same inbred line. For all primer pairs, over 99% of the reads aligned to the sequences representing the ten maize chromosomes, and 27–57% of the reads were aligned with no mismatches to a single locus (Table 4). Nearly 40% of the reads did not match perfectly to a genomic location, and the vast majority of these are likely to be the result of sequencing errors and incomplete genome. At this early stage of the bioinformatics, we have chosen to disregard these tags, but they could obviously be incorporated in most sophisticated pipelines.

**Table 4.** Read alignments for sample B73

| Primer Pair | Number of Reads | Read Alignment (% of Reads)* |  |  |  | Genomic Loci** |  |  |  |
| --- | --- | --- | --- | --- | --- | --- | --- | --- | --- |
|  |  | Aligned | Primary Chromosomes | Perfect Match | Unambiguous | Distinct Loci | Depth ≥ 10 Reads | Interlocus Distance (Kb) |  |
|  |  |  |  |  |  |  |  | Mean | Median |
| 1.2 | 165,505 | 99.99 | 99.85 | 61.14 | 43.08 | 7,489 | 3,225 | 142 | 96 |
| 2.4 | 164,242 | 100 | 99.55 | 63.10 | 56.96 | 2,422 | 1,901 | 424 | 286 |
| 3.1 | 147,336 | 99.99 | 99.76 | 60.49 | 33.89 | 5,610 | 1,891 | 186 | 125 |
| 4.1 | 172,066 | 100 | 99.81 | 57.99 | 26.58 | 1,573 | 1,353 | 670 | 438 |

\* Order of filters is left to right; only reads passing all preceding filters are evaluated for the percentage calculation.
\*\* Order of filters is left to right; only loci passing all preceding filters are evaluated for locus counts.

For the sequences with perfect matches, 46% (primer pair 4.1) to 90% (primer pair 2.4) were in unambiguous locations in the genome. The in silico primer selection algorithm had predicted that we would have had 80% (primer pair 4.1) to 95% (primer pair 2.4) uniqueness, so while the bioinformatics was a useful guide, it is clear that some primers are either amplifying non-target sequences or the reference genome is missing substantial numbers of copies for certain elements. Figure 3 clearly shows a secondary lower depth peak for primers pairs 4.1 and 3.1 suggesting off-target amplification. Primer pair 4.1 also has a long high-depth tail, suggesting multiple identical loci. Exact coordinates for all loci are provided in supplemental files S3–S6, and a summary of locus distribution across the maize genome and gene space for primer pair 2.4 is presented in Figure 2.

**Figure 2.**
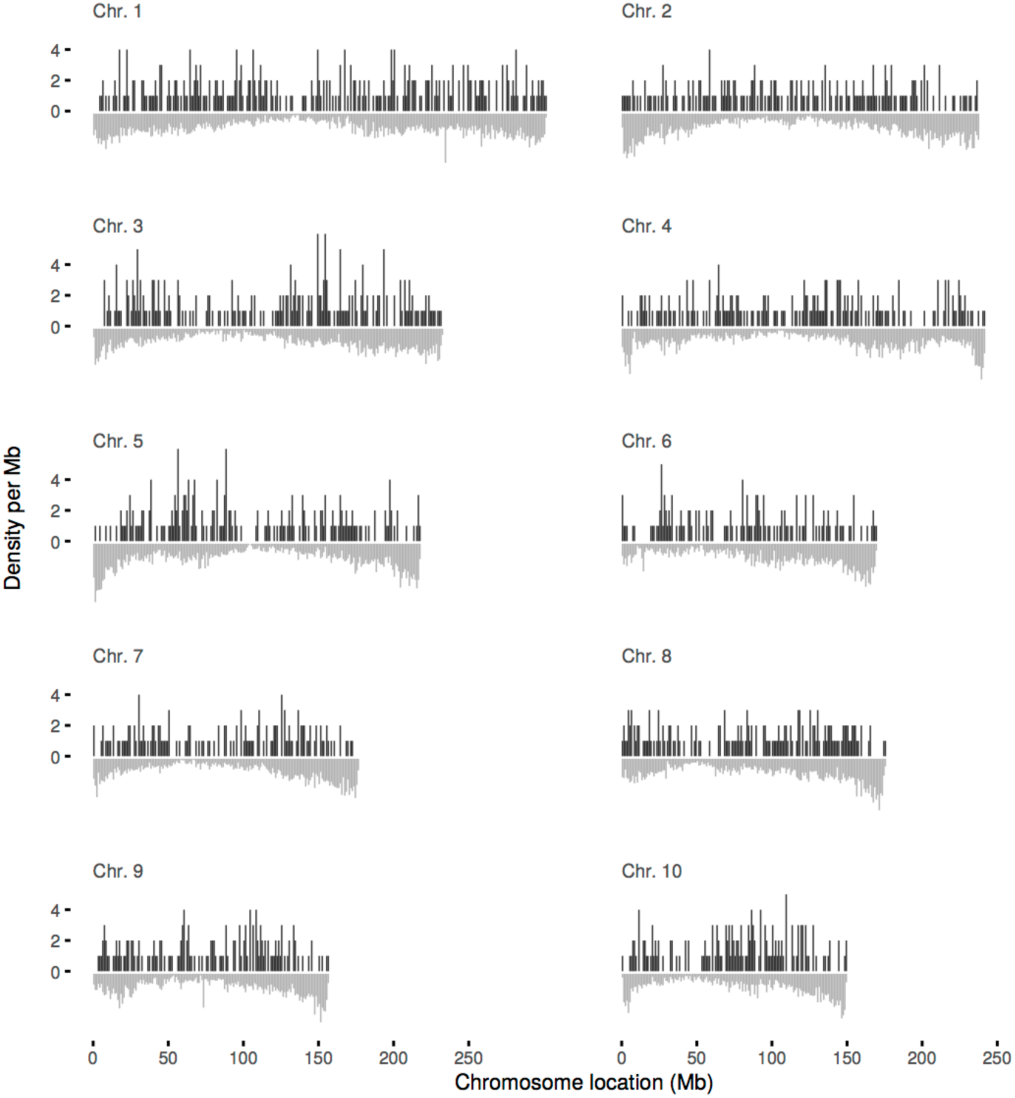
Alignment locus distribution across the maize genome and gene space. Alignment locus counts per Mb are presented as upwards black vertical bars with longer bars representing higher locus density, and range between 1 and 5 loci per Mb. Normalized gene model density is presented as downwards gray vertical bars, with longer bars representing higher density. Only results for primer pair 2.4 in sample B73 are presented.

**Figure 3.**
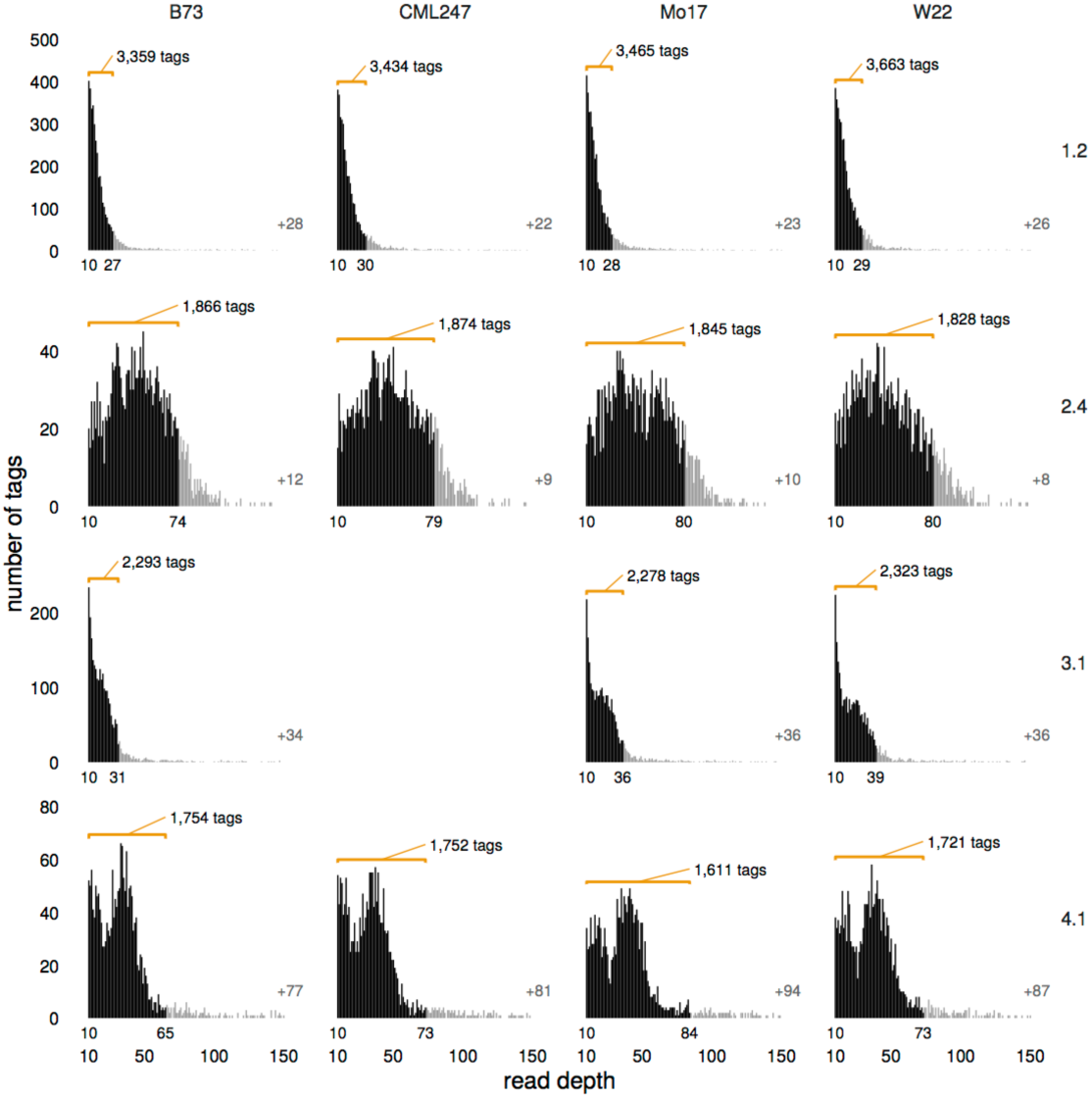
Sequence tag depth distribution in four deeply sequenced samples. For each sequence tag, read depth represents the number of reads with that exact sequence after quality filtering. The number of tags with read depth exceeding 150 reads is indicated by a gray number in the lower right margin of each distribution. The subset of tags selected for analysis is highlighted in black and indicated by a labeled orange bracket. The exact read depth cutoffs are indicated below each distribution along the x axis. No data are available for primer pair 3.1 in sample CML247.

#### Tag-based analysis Primer pair evaluation

The processed reads were further trimmed to the longest informative sequence expected for a given primer pair (130 bp for primer pairs 1.2, 2.4, and 4.1; 120 bp for primer pair 3.1), and only full length reads were retained for further analysis. For each sample and primer pair, the reads were collapsed into distinct sequences (sequence tags) and the depth of sequencing for each tag was recorded. We evaluated the tags’ read depth distribution for each of the four deeply sequenced samples (B73, CML247, W22, Mo17), and selected primer pair 2.4 for further analysis (see Discussion). Sequence tags were further filtered based on read depth. Any tags with a read depth less than 10 were discarded in order to remove likely sequencing and PCR artefacts, and the 10% most deeply sequenced tags were discarded in order to remove likely multi-locus tags (Figure 3).

#### Pairwise divergence estimates

Pairwise similarity between sequence tags from the B73 sample was evaluated using the k-tuple measure [26] as implemented in the Clustal Omega software (version 1.2.1; [27]) and results for primer pair 2.4 are presented in Figure 4. This highlights that while repetitive elements are similar they are far from identical with an average of 10% divergence.

**Figure 4.**
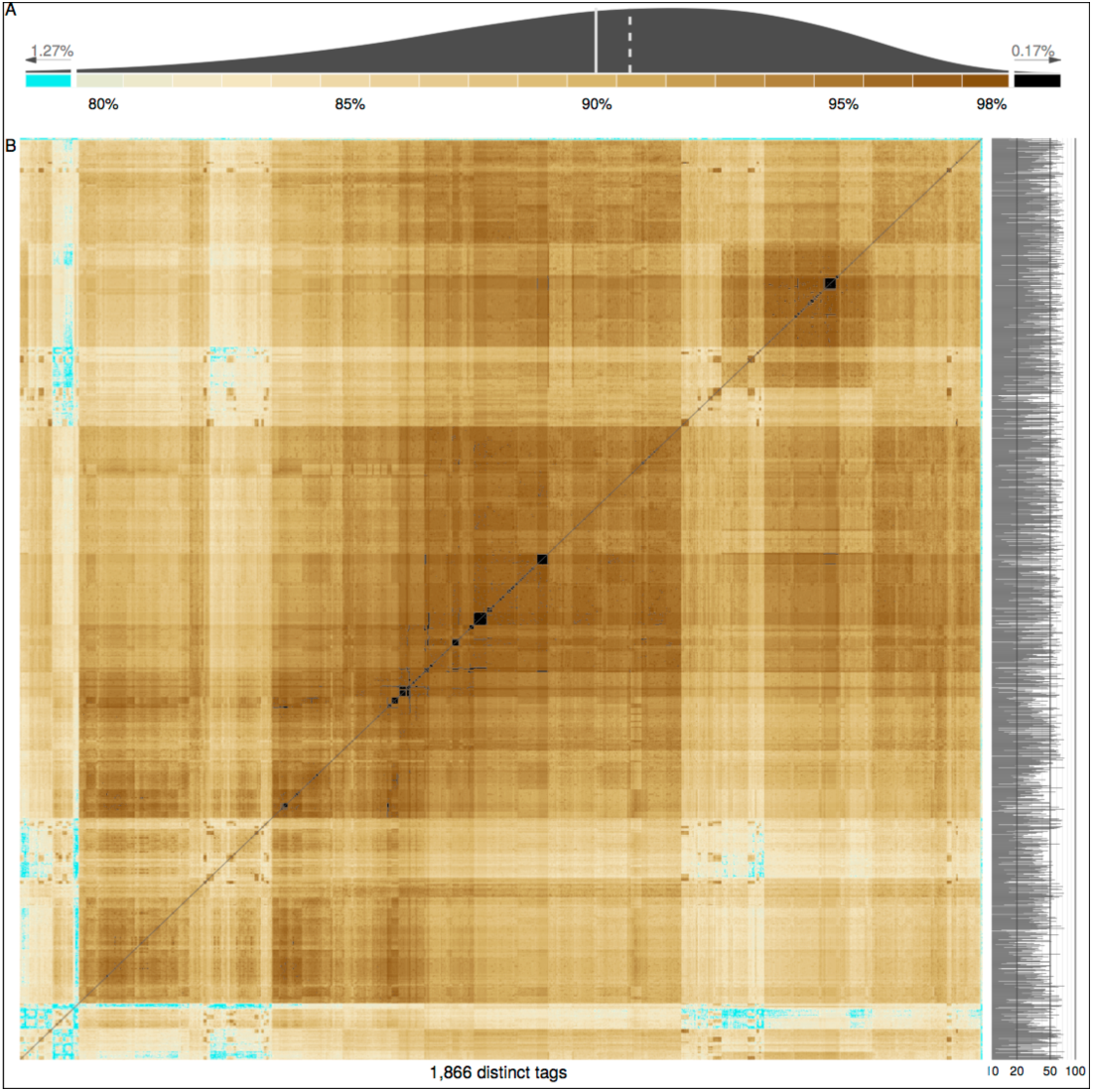
Pairwise similarity of sequence tag markers. Overall distribution of pairwise nucleotide similarity between tags (A) and heatmap of all pairwise sequence tag comparison results (B) from primer pair 2.4 in B73. Pairwise comparisons with less than 80% similarity (25 or more bp different; 1.27% of all comparisons) are highlighted in cyan. Pairwise comparisons with more than 98% similarity (1 bp different; 0.17 % of all comparisons) are highlighted in black. In A, vertical lines indicate the mean (solid) and median (dashed) values, and the colored blocks along the x axis represent the heatmap color scale. In B, the read depth for each tag is presented in the right margin on a log scale.

#### Genotyping of recombinant inbred population

Sequence tags amplified with primer pair 2.4 from the B73 and CML247 samples, representing parental genotypes of the recombinant inbred population, were assessed for their frequency in the recombinant inbred population with the expectation that each parent’s tags would be present, on average, in 50% of the progeny. The tag frequency data was modelled as a mixture of at most two one dimensional Gaussian distributions with variable variance using the package Mclust (version 5.2; https://cran.r-project.org/web/packages/mclust; [28]) for the R Statistical Computing Environment (version 3.2.2; https://www.R-project.org; [29]). For each sample, the Gaussian distribution with the majority contribution was selected and values delimiting two standard deviations around the mean were used as the lower and upper limits of tag frequency to further filter the sequence tags. Finally, only sequence tags specific to each parental genotype were kept for tag-based genotyping.

In order to precisely anchor the tags thus selected from the B73 sample on the physical map, they were re-aligned against the maize reference genome using bwa-mem with the following parameters: -M -c 100000 -L 500,500 -T 0. Tags having primary alignments (SAMtools flag -F 0xf0c) with n mismatches on the ten maize chromosomes were retained and further filtered to remove those with ambiguous alignments, define as having a secondary alignment with the same alignment score as the primary alignment. The recombinant inbred population was genotyped using these B73 tags as dominant markers, with genotype calls supported by less than 2 reads marked as unknown. These genotype calls were compared to the subset of GBS-based genotype calls from the Glaubitz et al. [30] data set (ZeaGBSv2.7;http://cbsusrv04.tc.cornell.edu/users/panzea/download.aspx?filegroupid=4) corresponding to the same accessions (Figure 5).

**Figure 5.**
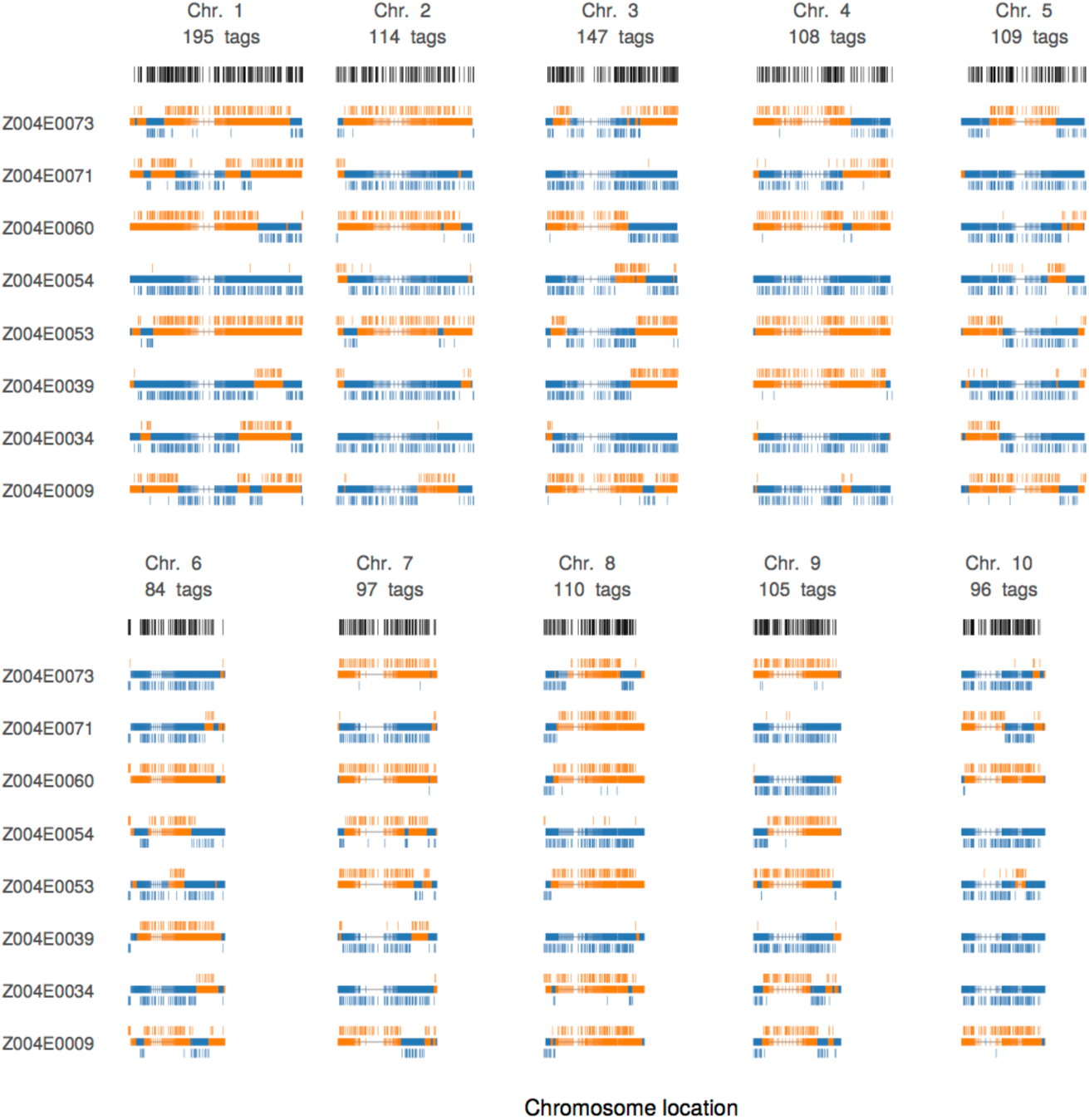
Recombination block identification in a recombinant inbred population Genotype assignments along the 10 maize chromosomes are shown for a subset of 8 samples, with blue indicating the B73 genotype and orange indicating the CML247 genotype. For each chromosome by sample plot, the central horizontal bar represents genotype assignments inferred from the dense GBS-based genotyping assay. The genotype assignments inferred from the rAmpSeq data are presented above (CML247 genotypes) and below (B73 genotypes). Below each chromosome label, the number of tags aligned to that chromosome and their physical location are indicated in gray text and black dashes respectively. Both GBS and rAmpSeq markers are presented as.

### Discussion

The great advantage of rAmpSeq is its simplicity. The method takes advantage of today’s longer, high-quality SBS reads and multiplexing strategies in combination with a simple PCR amplification. This simplicity results in some great benefits - the cost is extremely low and DNA preps are simple. The biggest side benefit from targeting repetitive sequences is that reduced PCR amplicon competition results in relatively even amplification profiles. Targeting repetitive sequences works well because of the diverse biology and ecology of retrotransposons that exist within genomes. Today, a protocol like rAmpSeq is reasonable because ignoring 20-50% of data is cost-effective for genotyping with current next-generation sequencing throughput.

Maize, like most plants, has a tremendous diversity of retrotransposons [31,32]. It is not widely recognized that, although transposons are repetitive, their sequences are not identical. Analysis of divergence between various amplicons shows that many families that share stretches of nearly identical sequence can have intervening regions with 10-15% divergence from most other amplicons (Figure X). Therefore, sequencing reads of 150 nucleotides can provide more than 15 diagnostic variants to differentiate paralogs.

There are still a number of limitations of rAmpSeq bioinformatics that can be greatly improved. The current approach treats each sequence tag as a unique dominant marker. This approach, however, does not capture the rich evolutionary history present in these sequences. Reconstructing an evolutionary network is relatively straightforward, but because of the substantial sequence depths required, PCR and sequencing errors also need to be modeled across the network. The UNEAK pipeline [33], used for calling GBS SNPs in species that lack a genome sequence, builds many small networks and identifies a subset of simple allelic pairs that are clearly above the sequencing error rate. Because rAmpSeq is focused at a much smaller portion of the genome with much higher coverage, PCR and sequencing error rates can be modeled for every nucleotide, and can be used to rigorously differentiate homologs from errors.

The dominant markers identified by the rAmpSeq sequence analysis presented here can also be converted to co-dominant markers through several strategies. First, as mentioned above, sequence networks can be used to differentiate allelic divergence from errors. Second, depth of coverage at a locus can be a reliable signal for presence/absence. Third, in situations where mapping populations exist, genetics can be used to used to identify putative alleles that are then confirmed by sequence divergence. Linkage disequilibrium mapping can also be used resolve the locations of tags in a similar manner [34]. Finally, there are many tags that map so closely (<<1cM) that they essentially function as genetic alleles, even though detailed structural genomics suggests they are not at the same position. Cursory analyses with the above dataset indicated that this situation was common, and nearly all dominant alleles could be converted to co-dominant suites of alleles. Future work will formally convert these features into a probabilistic model.

Most imputation approaches currently rely on using co-dominant markers. There is obviously substantial information content variation originating from different classes of repetitive sequences. For the maize samples analyzed here, it appears that some of the older families of repetitive sequences (Primer 9) produced the most valuable profiles, both for mapping SNPs and presence/absence variation. In lower diversity species, however, maximizing presence/absence variation may be most important, and, hence, targeting relatively young retrotransposons would be most effective. The vast majority of higher eukaryotes with genomes larger than 500 Mbp are likely to have plenty of repetitive families with genome-wide distributions. In these cases, it would be quite possible to multiplex to 2 or more repetitive families in a single reaction or use degenerate PCR primers to expand the amplification target range. While this strategy would take some optimization to reduce competition between families, it would provide a straightforward approach to providing enough markers in small genomes.

The cost profile of rAmpSeq is very competitive with other approaches. At 3000-plex, rAmpSeq would provide an estimated estimate $2.30 per sample (Table 5).

**Table 5.** Estimated costs for genotyping at 3000-plex.

| Process | Cost/Sample at 3000-plex |
| --- | --- |
| DNA extraction (CTAB) | \$ 0.25 |
| rAmpSeq PCR and sample cleanup | \$ 0.55 |
| Illumina dual index PCR and sample cleanup | \$ 0.60 |
| DNA sequencing (NextSeq 500, 1X150) | \$ 0.90 |
| Total | \$ 2.30 |

The greatest disadvantage of the protocol are that repetitive sequences are more difficult to analyze bioinformatically. Both the initial anchoring of sequences to the genome and differentiating paralogs from orthologs are particularly challenging. In the breeding context, where large numbers of segregants are scored from a cross, these issues can all be sorted out robustly using genetics. In conservation biology, sampling of the offspring from a few key mothers or fathers of the species of interest prior to genotyping entire populations could provide the clear linkage signal necessary to differentiate and to anchor novel paralogs. While the bioinformatics can be challenging, the TASSEL GBS pipeline provides a model of how to capture the species knowledge in a discovery run and database, which can then be shared across the species and further researchers only use a single step production run [30].

While the bioinformatic resources are not yet fully developed, rAmpSeq already has tremendous value in species with quality reference genomes. rAmpSeq also has the potential to be integrated with other amplicon-based sequencing approaches. For example, in many breeding contexts 10 to 20 functional loci are often genotyped regularly to ensure selection for disease resistance or quality traits. Combining these assays with rAmpSeq can provide the background markers that are needed for genome-wide predictions and genomic selection. The primer titration necessary to optimize PCR would require some effort, but this approach would help provide unambiguous results at key functional loci and near complete coverage of segregation for the rest of the genome.

The last decade saw genomics used in scientific discovery for thousands of species, but breeding or conservation applications were strongly felt only for a few dozen species. Genomic technologies, like rAmpSeq, that increase ease and reduce the cost of genotyping to the price of a cheap cup of coffee, help take the applied genomics revolution to all corners of the earth. Potential applications could be as diverse as breeding yams in Africa, understanding the migration of butterflies, and conservation and adaptation of trees to climate change.

## Acknowledgements

We thank Cornell’s BRC Genomics Facility for DNA sequencing. This work was supported by USDA-ARS and National Science Foundation Grants IOS-1238014.

## References

1. Meuwissen TH, Hayes BJ, Goddard ME. Prediction of total genetic value using genome-wide dense marker maps. Genetics. 2001;157: 1819–1829.

2. Varshney RK, Ribaut J-M, Buckler ES, Tuberosa R, Rafalski JA, Langridge P. Can genomics boost productivity of orphan crops? Nat Biotechnol. 2012;30: 1172–1176.

3. Altshuler D, Pollara VJ, Cowles CR, Van Etten WJ, Baldwin J, Linton L, et al. An SNP map of the human genome generated by reduced representation shotgun sequencing. Nature. 2000;407: 513–516.

4. Elshire RJ, Glaubitz JC, Sun Q, Poland JA, Kawamoto K, Buckler ES, et al. A robust, simple genotyping-by-sequencing (GBS) approach for high diversity species. PLoS One. dx.plos.org; 2011;6: e19379.

5. Gore MA, Chia J-M, Elshire RJ, Sun Q, Ersoz ES, Hurwitz BL, et al. A first-generation haplotype map of maize. Science. 2009;326: 1115–1117.

6. Van Tassell CP, Smith TPL, Matukumalli LK, Taylor JF, Schnabel RD, Lawley CT, et al. SNP discovery and allele frequency estimation by deep sequencing of reduced representation libraries. Nat Methods. Nature Publishing Group; 2008;5: 247–252.

7. Liu CL, Schreiber SL, Bernstein BE. Development and validation of a T7 based linear amplification for genomic DNA. BMC Genomics. 2003;4: 19.

8. Yang S, Fresnedo-Ramírez J, Wang M, Cote L, Schweitzer P, Barba P, et al. A next-generation marker genotyping platform (AmpSeq) in heterozygous crops: a case study for marker-assisted selection in grapevine. Hortic Res. 2016;3: 16002.

9. Saghai-Maroof MA, Soliman KM, Jorgensen RA, Allard RW. Ribosomal DNA spacer-length polymorphisms in barley: mendelian inheritance, chromosomal location, and population dynamics. Proc Natl Acad Sci U S A. 1984;81: 8014–8018.

10. Fritz GN, Conn J, Cockburn A, Seawright J. Sequence analysis of the ribosomal DNA internal transcribed spacer 2 from populations of Anopheles nuneztovari (Diptera: Culicidae). Mol Biol Evol. 1994;11: 406–416.

11. Buckler ES 4th, Holtsford TP. Zea systematics: ribosomal ITS evidence. Mol Biol Evol. 1996;13: 612–622.

12. Buckler ES 4th, Ippolito A, Holtsford TP. The evolution of ribosomal DNA: divergent paralogues and phylogenetic implications. Genetics. 1997;145: 821–832.

13. Gyllensten U, Sundvall M, Ezcurra I, Erlich HA. Genetic diversity at class II DRB loci of the primate MHC. J Immunol. 1991;146: 4368–4376.

14. Purugganan MD, Wessler SR. Transposon signatures: species-specific molecular markers that utilize a class of multiple-copy nuclear DNA. Mol Ecol. 1995;4: 265–269.

15. Casa AM, Brouwer C, Nagel A, Wang L, Zhang Q, Kresovich S, et al. The MITE family heartbreaker (Hbr): molecular markers in maize. Proc Natl Acad Sci U S A. 2000;97: 10083–10089.

16. Chang R-Y, O’Donoughue LS, Bureau TE. Inter-MITE polymorphisms (IMP): a high throughput transposon-based genome mapping and fingerprinting approach. Theor Appl Genet. Springer-Verlag; 2001;102: 773–781.

17. McMullen MD, Kresovich S, Villeda HS, Bradbury P, Li H, Sun Q, et al. Genetic properties of the maize nested association mapping population. Science. sciencemag.org; 2009;325: 737–740.

18. Rogers SO, Bendich AJ. Extraction of total cellular DNA from plants, algae and fungi. In: Gelvin SB, Schilperoort RA, editors. Plant Molecular Biology Manual. Springer Netherlands; 1994. pp. 183–190.

19. Bradbury PJ, Zhang Z, Kroon DE, Casstevens TM, Ramdoss Y, Buckler ES. TASSEL: software for association mapping of complex traits in diverse samples. Bioinformatics. 2007;23: 2633–2635.

20. Untergasser A, Cutcutache I, Koressaar T, Ye J, Faircloth BC, Remm M, et al. Primer3–new capabilities and interfaces. Nucleic Acids Res. 2012;40: e115.

21. Martin M. Cutadapt removes adapter sequences from high-throughput sequencing reads. EMBnet.journal. 2011;17: 10–12.

22. Kersey PJ, Allen JE, Armean I, Boddu S, Bolt BJ, Carvalho-Silva D, et al. Ensembl Genomes 2016: more genomes, more complexity. Nucleic Acids Res. 2016;44: D574–D580.

23. Li H. Aligning sequence reads, clone sequences and assembly contigs with BWA-MEM. arXiv:13033997 [q-bio]. 2013; Available: http://arxiv.org/abs/1303.3997

24. Li H, Handsaker B, Wysoker A, Fennell T, Ruan J, Homer N, et al. The Sequence Alignment/Map format and SAMtools. Bioinformatics. 2009;25: 2078–2079.

25. Quinlan AR, Hall IM. BEDTools: a flexible suite of utilities for comparing genomic features. Bioinformatics. 2010;26: 841–842.

26. Wilbur WJ, Lipman DJ. Rapid similarity searches of nucleic acid and protein data banks. Proc Natl Acad Sci U S A. 1983;80: 726–730.

27. Sievers F, Wilm A, Dineen D, Gibson TJ, Karplus K, Li W, et al. Fast, scalable generation of high-quality protein multiple sequence alignments using Clustal Omega. Mol Syst Biol. 2011;7: 539.

28. Fraley C, Raftery AE, Murphy TB, Scrucca L. Mclust Version 4 for R: Normal Mixture Modeling for Model-Based Clustering, Classification, and Density Estimation [Internet]. Department of Statistics, University of Washington; 2012. Report No.: 597. Available: https://www.stat.washington.edu/research/reports/2012/tr597.pdf

29. R Core Team. R: a language and environment for statistical computing. Vienna, Austria: R Foundation for Statistical Computing; 2015.

30. Glaubitz JC, Casstevens TM, Lu F, Harriman J, Elshire RJ, Sun Q, et al. TASSEL-GBS: a high capacity genotyping by sequencing analysis pipeline. PLoS One. 2014;9: e90346.

31. SanMiguel P, Gaut BS, Tikhonov A, Nakajima Y, Bennetzen JL. The paleontology of intergene retrotransposons of maize. Nat Genet. 1998;20: 43–45.

32. Baucom RS, Estill JC, Chaparro C, Upshaw N, Jogi A, Deragon J-M, et al. Exceptional diversity, non-random distribution, and rapid evolution of retroelements in the B73 maize genome. PLoS Genet. 2009;5: e1000732.

33. Lu F, Lipka AE, Glaubitz J, Elshire R, Cherney JH, Casler MD, et al. Switchgrass genomic diversity, ploidy, and evolution: novel insights from a network-based SNP discovery protocol. PLoS Genet. dx.plos.org; 2013;9: e1003215.

34. Lu F, Romay MC, Glaubitz JC, Bradbury PJ, Elshire RJ, Wang T, et al. High-resolution genetic mapping of maize pangenome sequence anchors. Nat Commun. Nature Research; 2015;6. doi:10.1038/ncomms7914

